# sLASER and PRESS Perform Similarly at Revealing Metabolite-Age Correlations

**DOI:** 10.1101/2023.01.18.524597

**Authors:** Steve C.N. Hui, Tao Gong, Helge J. Zöllner, Kathleen E. Hupfeld, Aaron T. Gudmundson, Saipavitra Murali-Manohar, Christopher W. Davies-Jenkins, Yulu Song, Yufan Chen, Georg Oeltzschner, Guangbin Wang, Richard A. E. Edden

**Affiliations:** The Russell H. Morgan Department of Radiology and Radiological Science, Johns Hopkins University School of Medicine, Baltimore, MD, USA; F.M. Kirby Research Center for Functional Brain Imaging, Kennedy Krieger Institute, Baltimore, MD, USA; Department of Radiology, Shandong Provincial Hospital, Cheeloo College of Medicine, Shandong University, Jinan, Shandong, China; Department of Radiology, Shandong Provincial Hospital Affiliated to Shandong First Medical University, Jinan, Shandong, China

**Keywords:** magnetic resonance spectroscopy, sLASER, PRESS, localization, aging

## Abstract

**Purpose:** To compare the respective ability of PRESS and sLASER to reveal biological relationships, using age as a validation covariate.

**Methods:** MRS data were acquired from 102 healthy volunteers using PRESS and sLASER in centrum semiovale (CSO) and posterior cingulate cortex (PCC) regions. Acquisition parameters included TR/TE 2000/30 ms; 96 transients; 2048 datapoints sampled at 2 kHz.

Spectra were analyzed using Osprey. Signal-to-noise ratio (SNR), full-width-half-maximum linewidth of tCr, and metabolite concentrations were extracted. A linear model was used to compare SNR and linewidth. Paired t-tests were used to assess differences in metabolite measurements between PRESS and sLASER. Correlations were used to evaluate the relationship between PRESS and sLASER metabolite estimates, as well as the strength of each metabolite-age relationship. Coefficients of variation were calculated to assess inter-subject variability in each metabolite measurement.

**Results:** SNR and linewidth were significantly higher (p<0.05) for sLASER than PRESS. Paired t-tests showed significant differences between PRESS and sLASER in most metabolite measurements. Metabolite measures were significantly correlated (p<0.05) for most metabolites between the two methods except GABA, Gln and Lac in CSO and GSH, Lac and NAAG in PCC. Metabolite-age relationships were consistently identified using both PRESS and sLASER. Similar CVs were observed for most metabolites.

**Conclusion:** The study results suggest strong agreement between PRESS and sLASER in identifying relationships between brain metabolites and age in CSO and PCC data acquired at 3T. sLASER is technically desirable due to the reduced chemical shift displacement artifact; however, PRESS performed similarly in ‘good’ brain regions at clinical field strength.

## 1 INTRODUCTION

Recent MRS community consensus (1,2) has recommended the sLASER (3) localization sequence over PRESS (4) for single-voxel ^1^H MRS. This recommendation is mainly motivated by the improved localization performance of sLASER, in particular its reduced chemical shift displacement artifact (CSDA) that results from higher slice-selection bandwidth. The CSDA occurs because metabolite signals with different Larmor frequencies experience the slice-selective RF pulses differently, resulting in a spatial displacement of the selected slice that is proportional to the bandwidth of the slice-selective pulse. Increased CSDA is undesirable for two main reasons: detecting different metabolite signals from slightly different locations makes the interpretation of results more challenging; and displacement of the water- and lipid-excited volume makes it more challenging to acquire high-quality data.

Phase-coherent refocusing – that is, the ability to apply a 180° pulse to transverse magnetization and yield a spin echo signal with the same phase throughout the selected slice – is more challenging than phase-coherent excitation, resulting in substantial CSDA in two of the three voxel dimensions. For example, commonly used sinc-Gaussian pulses have only one-third of the bandwidth for refocusing as for excitation at the same peak RF *B*_1,max_. For this reason, Philips and Canon scanners replace PRESS sinc-Gaussian refocusing pulses with an asymmetric amplitude-modulated waveform with about twice the bandwidth, and GE often runs PRESS with reduced refocusing flip-angle to increase bandwidth. sLASER avoids the difficulties of phase-coherent slice-selective refocusing by using a pair of mutually correcting inversion pulses, as described in greater detail below. The issue of CSDA worsens with increasing *B*_0_ field strength because the frequency dispersion of the spectrum increases. It has not generally been the case that the peak available *B*_1,max_ has scaled up linearly with *B*_0_ on higher-field scanners (due both to engineering and SAR constraints).

PRESS localization has been widely applied for over three decades (4,5). Test-retest coefficients of variation demonstrate reproducible results for multiple major metabolites at 1.5T and 3T (6-9). PRESS is a single-shot localization sequence consisting of one slice-selective 90° excitation pulse and two slice-selective 180° refocusing pulses, each being applied to select one of the three orthogonal directions for volume selection (4). The two refocusing pulses yield a spin echo that fully utilizes the available *M*_*z*_ magnetization to produce a full-intensity signal (10,11) for an SNR-efficient acquisition. The minimum achievable echo time (TE) for PRESS is typically 30-35 ms. CSDA can be reduced by replacing more traditional sinc-Gaussian-like pulses (as used in Siemens and GE PRESS sequences) with asymmetric ‘Murdoch’ pulses (as used in Philips and Canon PRESS sequences); however, the bandwidth of the RF pulses in PRESS is typically limited to 1–2 kHz due to peak *B*_1_ limitations (12) corresponding to 6-13% CSDA per ppm shift.

For a given B_1_ maximum, adiabatic full passage (AFP) 180° inversion pulses have much higher bandwidth than refocusing pulses, achieved by removing the constraint of phase coherence across the slice. For a refocusing pulse, spins throughout the selected slice experience the 180° rotation at the same time within the pulse, meaning that the majority of the ‘effort’ of the pulse is concentrated towards the middle (as most easily visualized in a sinc-Gaussian waveform). AFP pulses sweep across the slice inverting one side of the pulse early, and the other side late, distributing the ‘effort’ of the pulse more evenly throughout its duration and allowing for greater bandwidth with a given peak *B*_1_. This results in signals that are not phase-coherent across the slice. A pair of AFP pulses must therefore be used to define a slice, the pair being mutually refocusing and in combination yielding a signal that is phase-coherent across the slice, as in LASER (13) or sLASER (3,14-16). The resulting sLASER slice profile is therefore the square of the AFP inversion profile, a function that has rarely been plotted in the literature. Hyperbolic-secant AFP pulses have more recently been superseded by gradient-modulated RF pulses with bandwidth typically between 8-10 kHz including the bandwidth-modulated adiabatic selective saturation and inversion (BASSI) (17), frequency offset corrected inversion (FOCI) (18) and gradient_□modulated offset_□independent adiabaticity (GOIA) (19). Studies have demonstrated that sLASER has good reliability and reproducibility at a range of field strengths (15,20,21). sLASER implemented with gradient-modulated RF pulses retains the single-shot full-intensity signal and can generally be acquired with TEs only slightly longer than PRESS (given similar crusher gradient areas) and substantially reduced CSDA (1.3% per ppm for 10-kHz bandwidth). The optimal crusher gradient setup and pulse order for sLASER are still under debate (3,15,16,22,23). Although sLASER sequences are available as product or research sequences for all major vendors, access to these sequences is less straightforward than for PRESS which has long-standing product status.

There are not many studies directly comparing the performance of PRESS and sLASER, especially using large-cohort in vivo data. Most prior studies have compared methodological aspects in small sample sizes. One early study compared sinc-180°, AFP, and FOCI pulses with bandwidths of 1.45, 2.5 and 12.5 kHz respectively in terms of B_1_ inhomogeneity, localization accuracy, and signal recovery and demonstrated that experimental CSDA matched predictions (24). Scheenen et al. also demonstrated a four-fold reduction in CSDA from sLASER using AFP pulses compared to PRESS using Mao-180° pulses (14). Another study found significant differences in metabolite concentrations between two protocols using PRESS and sLASER and different shimming methods at 3T (16). They reported a CSDA of 11.6% per ppm for PRESS, and 2.0% per ppm for sLASER. Water linewidths were 10.5 Hz and 6.1 Hz in PRESS and sLASER respectively, making it difficult to separate the benefit of sLASER localization from the benefits of improved shimming. A recent study tested CSDA in vitro using a phantom containing fat and water compartments and data reproducibility on myocardial fatty acid and creatine in vivo using PRESS and sLASER as well as STEAM (25). Their results indicated CSDA of 28% and 10% for PRESS and sLASER, respectively (25). Most comparison studies between PRESS and sLASER have focused on the technical aspect of CSDA and tested whether practical results matched those calculated theoretically. However, there is a lack of large-scale in vivo experimental studies showing whether improved localization and CSDA translate into improved metabolite quantification, which is the ultimate goal.

While excellent localization is desirable, localization is not the single factor limiting single-voxel MRS, and it is important not only to demonstrate that sLASER is theoretically preferable, but also that the newer methodology has greater power to interrogate in vivo biochemistry. There is an extensive literature documenting age-related changes in metabolite levels measured with MRS (26-29), making age an appropriate external validation variable to compare the performance of PRESS and sLASER. The aim of this study was to compare in vivo brain metabolite concentration measurements and spectral quality between the two sequences, using the same shimming approach. If sLASER is a better methodology for applied MRS studies of the brain than PRESS, we would expect it to show: increased SNR and reduced linewidth due to improved localization; decreased variance in metabolite measurements; and increased sensitivity to metabolite-age correlations.

## 2 METHODS

### Acquisition

A sex- and age-balanced cohort of 102 healthy volunteers was recruited with local IRB approval (53 female; aged 20-69; mean 44.7 ± 13.3 years). Exclusion criteria included contraindications for MRI and a history of neurological and psychiatric illness. A Philips 3T MRI scanner (Ingenia CX, Philips Healthcare, The Netherlands) with a 32-channel phased-array head coil for RF receive was used to acquire MRS data with PRESS and sLASER. Voxels were localized in the predominantly white matter centrum semiovale (CSO) and the predominantly gray matter posterior cingulate cortex (PCC) regions (30 × 26 × 26 mm^3^) as shown in Figure 1. T_1_-weighted MPRAGE (TR/TE/ 6.9/3.2 ms; FA 8°; 1 mm^3^ isotropic resolution) was acquired for voxel positioning and tissue segmentation. PRESS localization employed the 180° ‘Murdoch’ amplitude-modulated refocusing pulses (bandwidth 1.3 kHz; duration 6.90 ms, max. B_1_: 13.5µT) and sLASER employed the gradient-modulated GOIA-WURST pulses (bandwidth 10 kHz; duration 4.5 ms, max. B_1_: 15µT) (30). Both sequences were acquired with: TR/TE 2000/30 ms, 96 transients of 2048 datapoints sampled at 2 kHz with VAPOR (31) water suppression (bandwidth 140 Hz). A 20-mm slice-selective saturation pulse was applied to suppress subcutaneous lipid adjacent to the voxel. Water reference spectra were acquired without water suppression. The PRESS data in this manuscript have been previously analyzed in two studies (32,33).

**Figure 1.**
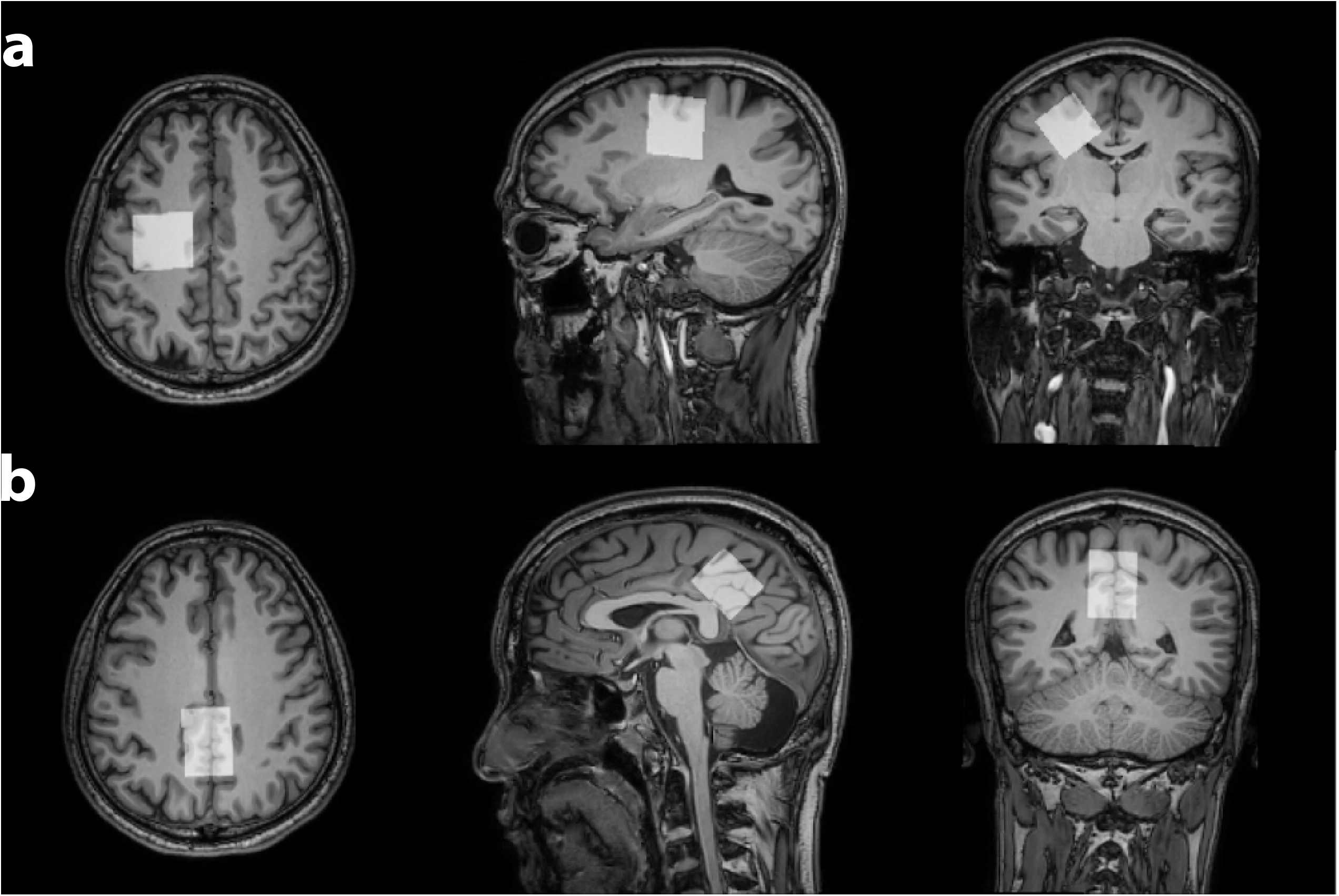
Data were acquired in 30 × 26 × 26 mm^3^ voxels in (a) left centrum semiovale (CSO) and (b) posterior cingulate cortex (PCC).

### Quantification

Spectra were processed and modeled using the Osprey software v2.4 (34), following consensus recommendation for linear combination model fitting (35). PRESS and sLASER basis sets consisting of 18 simulated metabolite basis functions and sequence-specific cohort-averaged (33) measured mobile macromolecule (MM) functions were employed. The metabolite basis set was generated from a fully localized 2D density-matrix simulation (101 × 101 resolution) carried out across a field of view that extends 50% larger than the nominal voxel size in both dimensions using real vendor pulse waveforms and sequence timings (36,37). Metabolites included in the basis set were ascorbate Asc; aspartate Asp; creatine Cr; negative creatine methylene -CrCH2; gamma-aminobutyric acid GABA; glycerophosphocholine GPC; glutamine Gln; glutamate Glu; glutathione GSH; lactate Lac; myo-inositol mI; N-acetylaspartate NAA; N-acetylaspartylglutamate NAAG; phosphocholine PCh; phosphocreatine PCr; phosphoethanolamine PE; scyllo-inositol sI; and taurine Tau. Cohort-mean measured sequence-specific MM basis functions were derived as described in a previous study that investigated the age trajectory of MM signals in the spectrum using the same dataset (32). Briefly, individual-subject ‘clean’ MM spectra were modeled with a flexible spline of 0.1 ppm knot spacing and the overall mean ‘clean’ MM spectra were generated and employed as the MM basis functions (33). The basis set—including simulated metabolite and measured MM basis functions—was incorporated into the Osprey software for modeling.

Brain tissue segmentation was performed using the Osprey-integrated SPM12 (38) to yield relative tissue volume fractions of gray matter, white matter, and cerebrospinal fluid for tissue correction. The water-reference data were quantified with a simulated water basis function in the frequency domain with a 6-parameter model (amplitude, zero-and first-order phase, Gaussian and Lorentzian line broadening, and frequency shift). Metabolite measurements were water-scaled and tissue-corrected for tissue-specific water visibility and relaxation times based on literature values (39). Data quality metrics (signal-to-noise ratio (SNR) and full-width-half-maximum (FWHM) linewidth of the 3-ppm tCr signal) were calculated for the processed metabolite spectra. SNR is defined in Osprey as the ratio of the tCr singlet peak height and the detrended standard deviation of the frequency-domain spectrum between -2 and 0 ppm. Linewidth was defined as the average of the peak-measured FWHM and the FWHM of a Lorentzian model of the tCr singlet. Visual inspection was performed to evaluate artifacts and contamination to ensure data quality according to consensus recommendations (1).

### Statistical analysis

All statistical analyses were performed using R v4.0.2 in RStudio v1.2.5019 (40). Metabolite concentrations from PRESS and sLASER data were measured for total NAA (tNAA=NAA+NAAG), total choline (tCho=GPC+PCh), total creatine (tCr=Cr+PCr), Glx (Glu+Gln) and individual contributions from Asp, GABA, Gln, Glu, GSH, Lac, mI, NAA, NAAG, PE and sI. Datapoints that had concentration measurement of 0 were excluded as they were interpreted as modeling failure. The Shapiro-Wilk test was used to test the normality of each variable. Two linear models were used to compare SNR and linewidth between PRESS and sLASER across both brain regions; these models included either tCr SNR or linewidth as the dependent variable, and sequence and voxel location as predictors. Paired t-tests were used to compare mean metabolite concentration measurements between PRESS and sLASER for normally distributed variables; otherwise, Wilcoxon signed-rank tests were used. Pearson correlation coefficients were used to test the relationship between PRESS and sLASER metabolite concentration measurements and correlations between age and each metabolite measurement; in cases of non-normally distributed variables, Spearman’s rank correlations were used. Coefficients of variation (CV) were calculated for each metabolite to compare the between-subject variation of metabolite measurements between PRESS and sLASER. P-values less than 0.05 were considered statistically significant.

## 3 RESULTS

Two PRESS and two sLASER spectra were excluded due to lipid contamination, presumably due to subject motion, and visible ethanol signal, based on visual inspection. Average metabolite and MM spectra with fit residual are shown in Figure 2. Average tCr SNRs were 97.5 ± 14.5 and 101.7 ± 17.1 and linewidths were 6.0 ± 0.7 Hz and 6.2 ± 1.2 Hz, respectively for PRESS and sLASER, indicating high data quality across both experiments and regions. SNR and linewidth fulfilled the minimum criterion for quality assessment suggested by the community consensus (1). The linear model indicated that SNR was significantly higher (p=0.004) by 4% for sLASER as shown in Figure 3a and linewidth was significantly higher (p=0.006) by 4.6% for sLASER as shown in Figure 3b. PCC had significantly higher (p<0.05) SNR and linewidth (105.2 ± 16.1 and 6.2 ± 0.8 Hz, respectively) than CSO (94.0 ± 13.7 and 5.99 ± 1.1 Hz).

**Figure 2.**
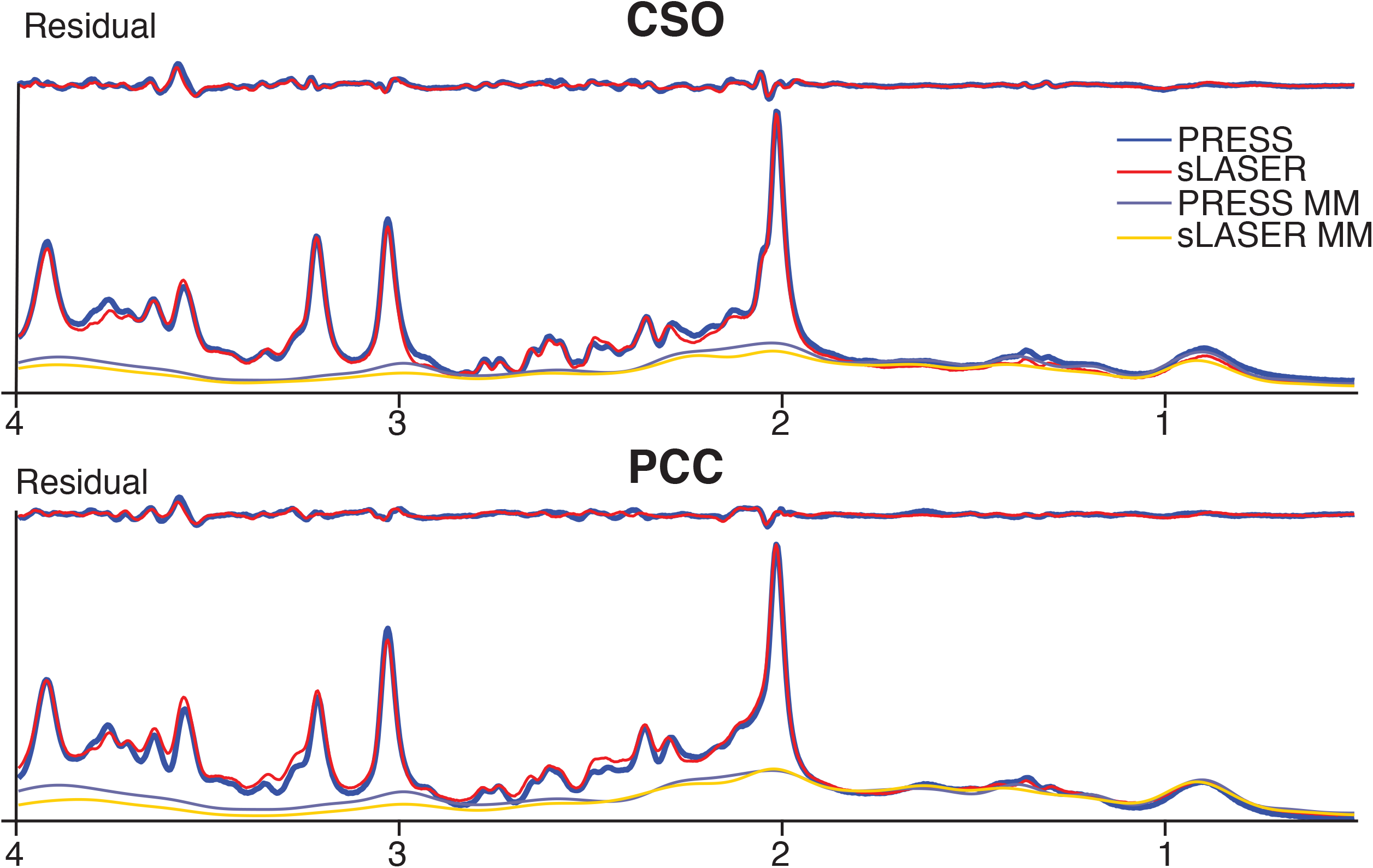
Cohort-mean spectra from CSO (above) and PCC (below). Mean metabolite spectra for PRESS and sLASER are overlaid. The mean LCM residuals (i.e. data – model) for each method and the mean modeled MM components are also shown.

**Figure 3.**
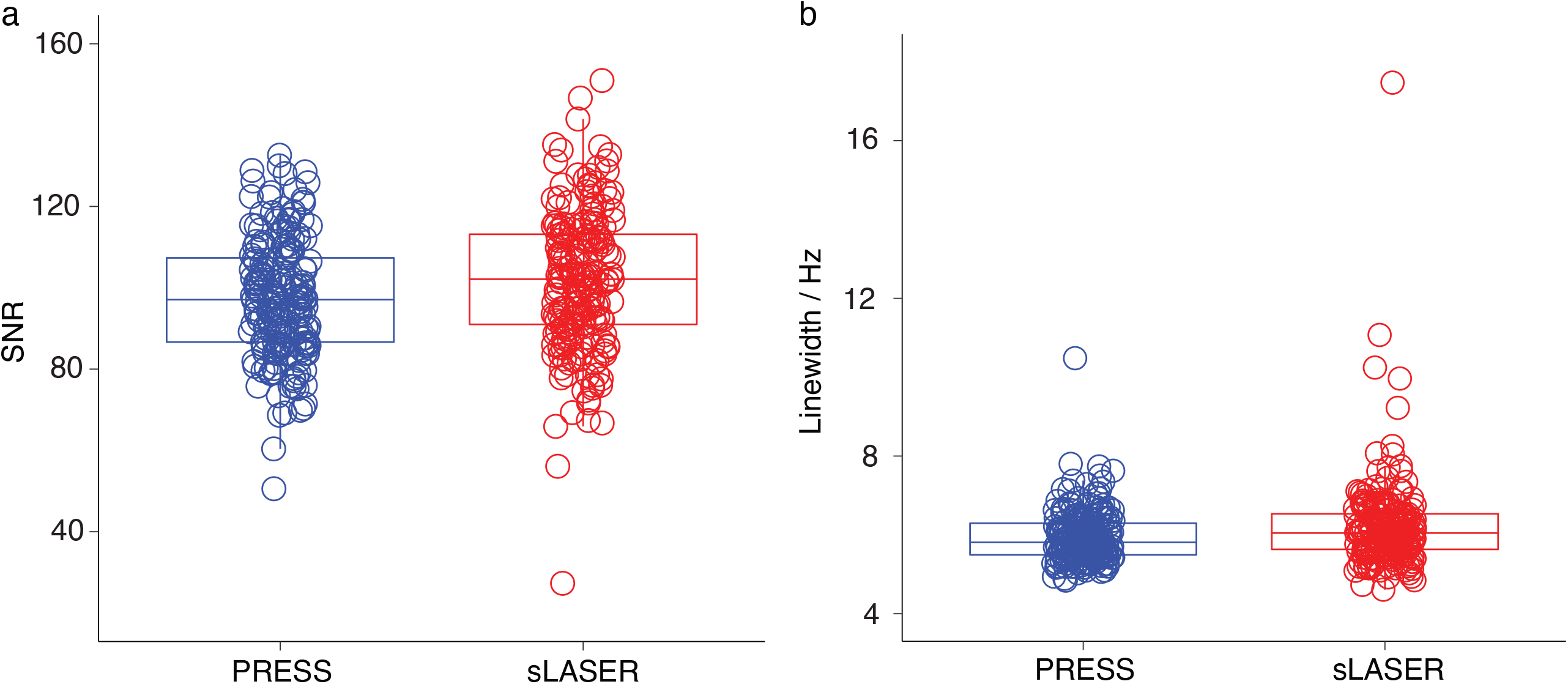
Box plots depicting (a) tCr SNR and (b) tCr linewidth, comparing PRESS and sLASER. Data acquired with PRESS and sLASER are depicted in dark blue and red, respectively.

Amplitude estimates of 0 were interpreted as modeling failure so were excluded from the correlation analysis; these were identified in 6 Gln fits, 8 Lac fits and 51 GABA fits in CSO and 16 Lac fits and 69 GABA fits in PCC. In CSO and PCC Shapiro-Wilk tests indicated that 4 (tCr, Gln, PE and GABA) out of 15 metabolites were non-normally distributed in CSO and 9 (tNAA, tCho, GABA, GSH, PE, NAA, tCr, Lac and mI) in PCC. Results suggested that metabolite concentration measurements were significantly different between PRESS and sLASER except for tNAA and sI in PCC as shown in Table 1. Statistically significant PRESS-sLASER correlations were observed for most modeled metabolites except GABA, Gln and Lac in CSO and GSH, Lac and NAAG in PCC as shown in Figure 4, but only with modest correlation coefficients in most cases (though up to 0.7 for tCho, tCr, and mI). In CSO, significant positive metabolite-age relationships were observed for both PRESS and sLASER for tCho, tCr, mI, and GSH as shown in Figure 5. In PCC, significant positive metabolite-age relationships were observed for both PRESS and sLASER for tCho, tCr and mI, and a negative age relationship was observed for Asp. PCC PRESS data suggested a positive correlation of age with GSH that sLASER did not and vice versa for Glu.

**Table 1.** Mean metabolite levels in institutional units and comparisons between PRESS and sLASER using paired t-test.

| CSO |  |  |  |  | PCC |  |  |  |  |
| --- | --- | --- | --- | --- | --- | --- | --- | --- | --- |
| Metabolite | n | PRESS<br>Mean (SD) | sLASER<br>Mean (SD) | *p* | Metabolite | n | PRESS<br>Mean (SD) | sLASER<br>Mean (SD) | *p* |
| tNAA | 100 | 14.91 (0.74) | 16.12 (0.77) | <0.001 | tNAA* | 100 | 15.16 (0.59) | 15.14 (0.79) | 0.52 |
| NAA | 100 | 12.60 (0.63) | 13.65 (0.60) | <0.001 | NAA* | 100 | 13.73 (0.62) | 13.57 (0.78) | <0.05 |
| NAAG | 100 | 2.28 (0.45) | 2.44 (0.45) | <0.001 | NAAG | 100 | 1.36 (0.34) | 1.50 (0.34) | <0.01 |
| tCho | 100 | 2.33 (0.23) | 2.27 (0.23) | <0.001 | tCho* | 100 | 1.95 (0.17) | 1.87 (0.23) | <0.001 |
| tCr* | 100 | 9.40 (0.65) | 9.68 (0.64) | <0.001 | tCr* | 100 | 12.28 (0.71) | 11.26 (0.88) | <0.001 |
| mI | 100 | 6.81 (1.02) | 7.10 (0.91) | <0.001 | mI* | 100 | 9.74 (1.03) | 8.43 (0.90) | <0.001 |
| Glx | 100 | 8.52 (1.24) | 11.76 (1.34) | <0.001 | Glx | 100 | 18.79 (1.83) | 17.45 (1.80) | <0.001 |
| Glu | 100 | 7.92 (0.89) | 9.62 (0.89) | <0.001 | Glu | 100 | 14.90 (1.11) | 13.20 (1.13) | <0.01 |
| Gln* | 94 | 0.67 (0.54) | 2.17 (0.62) | <0.001 | Gln | 100 | 3.94 (0.89) | 4.30 (0.91) | <0.01 |
| GSH | 100 | 1.95 (0.32) | 1.47 (0.24) | <0.001 | GSH* | 100 | 2.46 (0.31) | 1.77 (0.39) | <0.001 |
| Lac | 92 | 1.19 (0.46) | 1.41 (0.67) | <0.01 | Lac* | 84 | 1.01 (0.65) | 1.50 (1.00) | <0.001 |
| sI | 100 | 0.41 (0.14) | 0.53 (0.17) | <0.001 | sI | 100 | 0.50 (0.17) | 0.51 (0.18) | 0.387 |
| PE* | 100 | 4.95 (0.94) | 4.08 (0.91) | <0.001 | PE* | 100 | 3.96 (0.78) | 2.53 (0.78) | <0.001 |
| Asp | 100 | 2.29 (0.54) | 3.30 (0.46) | <0.001 | Asp | 100 | 4.11 (0.57) | 3.70 (0.53) | <0.01 |
| GABA* | 49 | 1.21 (0.60) | 0.26 (0.22) | <0.001 | GABA* | 31 | 1.67(0.71) | 0.16(0.20) | <0.001 |

\*Indicates that metabolite concentrations were non-normally distributed; Wilcoxon signed-rank test was used.

**Figure 4.**
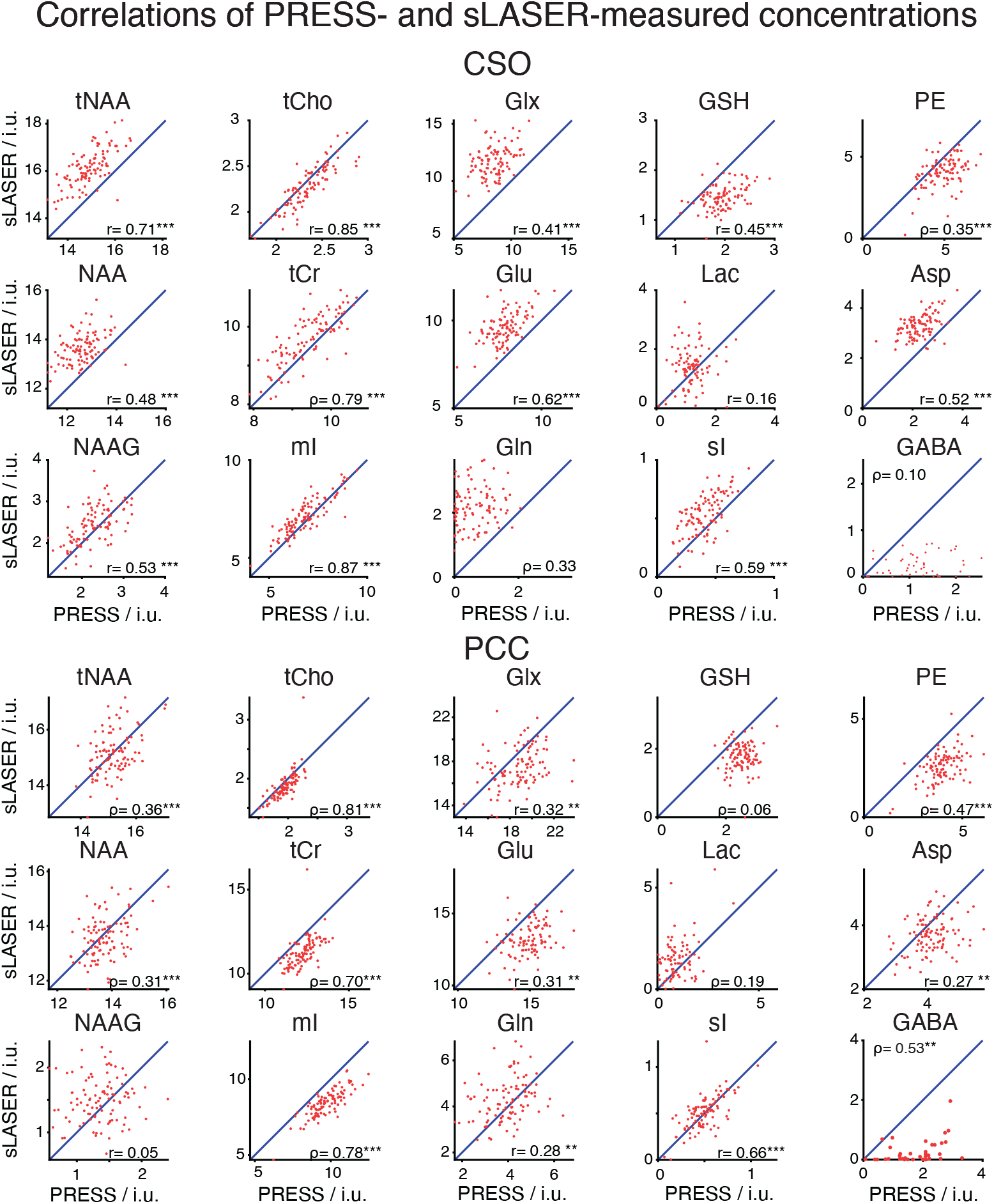
Correlations of PRESS- and sLASER-measured concentrations. Statistically significant correlations are indicated by asterisks (p<0.001***, p<0.01**, p<0.05*). The line indicates the x=y diagonal, not a modeled trendline.

**Figure 5.**
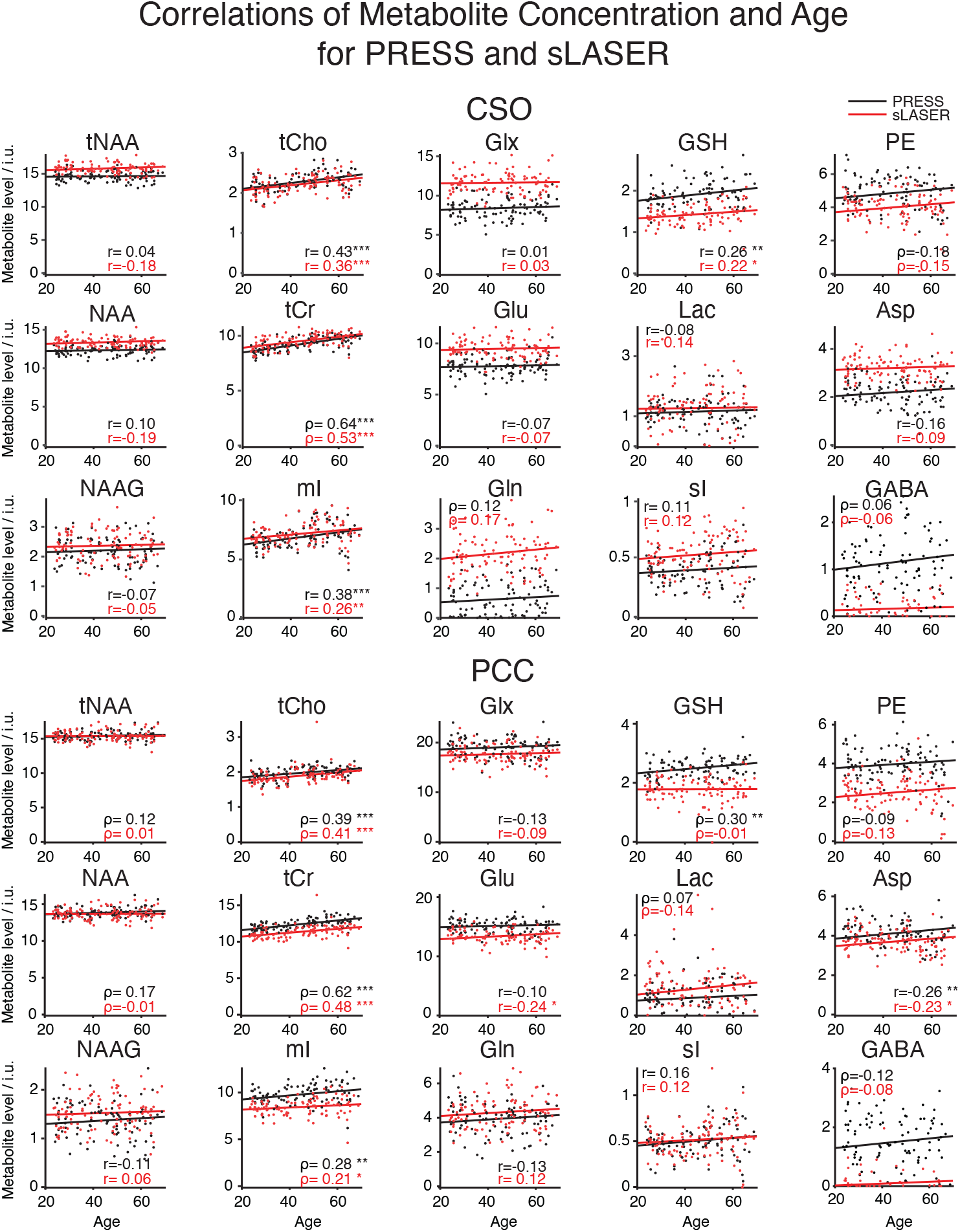
Correlations of metabolites concentration and age for PRESS (black) and sLASER (red). Linear trendlines are plotted for PRESS in black and sLASER in red. Statistically significant correlations are indicated by asterisks (p<0.001***, p<0.01**, p<0.05*).

Metabolite CVs for PRESS and sLASER are plotted on a logarithmic scale in Figure 6. Most metabolites of interest had similar CVs for PRESS and sLASER (ranging from 4-5% for NAA to over 30% for Lac, sI and GABA). PRESS had higher CVs for 10 out of 15 metabolites of interest (Asp, GSH, Gln, Glu, mI, NAA, NAAG, sI, tCr and tNAA) in CSO, and 4 out of 15 (Gln, mI, Lac, and NAAG) in PCC as shown in Table 2. GABA CVs showed high variability between PRESS (CSO:55%, PCC:50%) and sLASER (CSO:89%, PCC 124%); these were the highest CVs among all metabolites for both regions. Gln CVs also had high variability between PRESS (79%) and sLASER (30%) in the CSO region but not in PCC. CV differences between PRESS and sLASER were mostly below 10% for the other metabolites (majority were below 5% difference for both regions).

**Table 2.** Inter-subject coefficients of variation (%) for PRESS and sLASER in CSO and PCC.

|  | Asp | GABA | GSH | Gln | Glu | Glx | mI | Lac | NAA | NAAG | PE | sI | tCho | tCr | tNAA |
| --- | --- | --- | --- | --- | --- | --- | --- | --- | --- | --- | --- | --- | --- | --- | --- |
| CSO PRESS | 22.4 | 55.2 | 16.8 | 79.1 | 11.2 | 14.8 | 15.4 | 36.3 | 5.2 | 19.7 | 19.0 | 33.3 | 9.6 | 7.0 | 5.1 |
| CSO sLASER | 14.4 | 88.6 | 16.6 | 29.9 | 9.4 | 11.4 | 12.9 | 45.3 | 4.4 | 18.7 | 20.0 | 32.6 | 10.2 | 6.6 | 4.8 |
| *Difference* | *8.0* | *-33.4* | *0.2* | *49.3* | *1.8* | *-0.6* | *2.5* | *-9.1* | *0.8* | *1.0* | *-0.9* | *0.7* | *-0.6* | *0.4* | *0.3* |
| PCC PRESS | 13.8 | 50.4 | 12.1 | 23.0 | 7.5 | 9.8 | 10.0 | 58.3 | 4.2 | 24.8 | 17.3 | 31.9 | 8.7 | 5.7 | 3.5 |
| PCC sLASER | 14.2 | 123.6 | 19.6 | 21.5 | 8.1 | 10.3 | 9.6 | 56.2 | 5.7 | 22.6 | 27.6 | 33.1 | 9.5 | 6.4 | 5.2 |
| *Difference* | *-0.4* | *-73.2* | *-7.5* | *1.5* | *-0.6* | *-0.5* | *0.4* | *2.2* | *-1.5* | *2.2* | *-10.2* | *-1.1* | *-0.8* | *-0.7* | *-1.7* |

**Figure 6.**
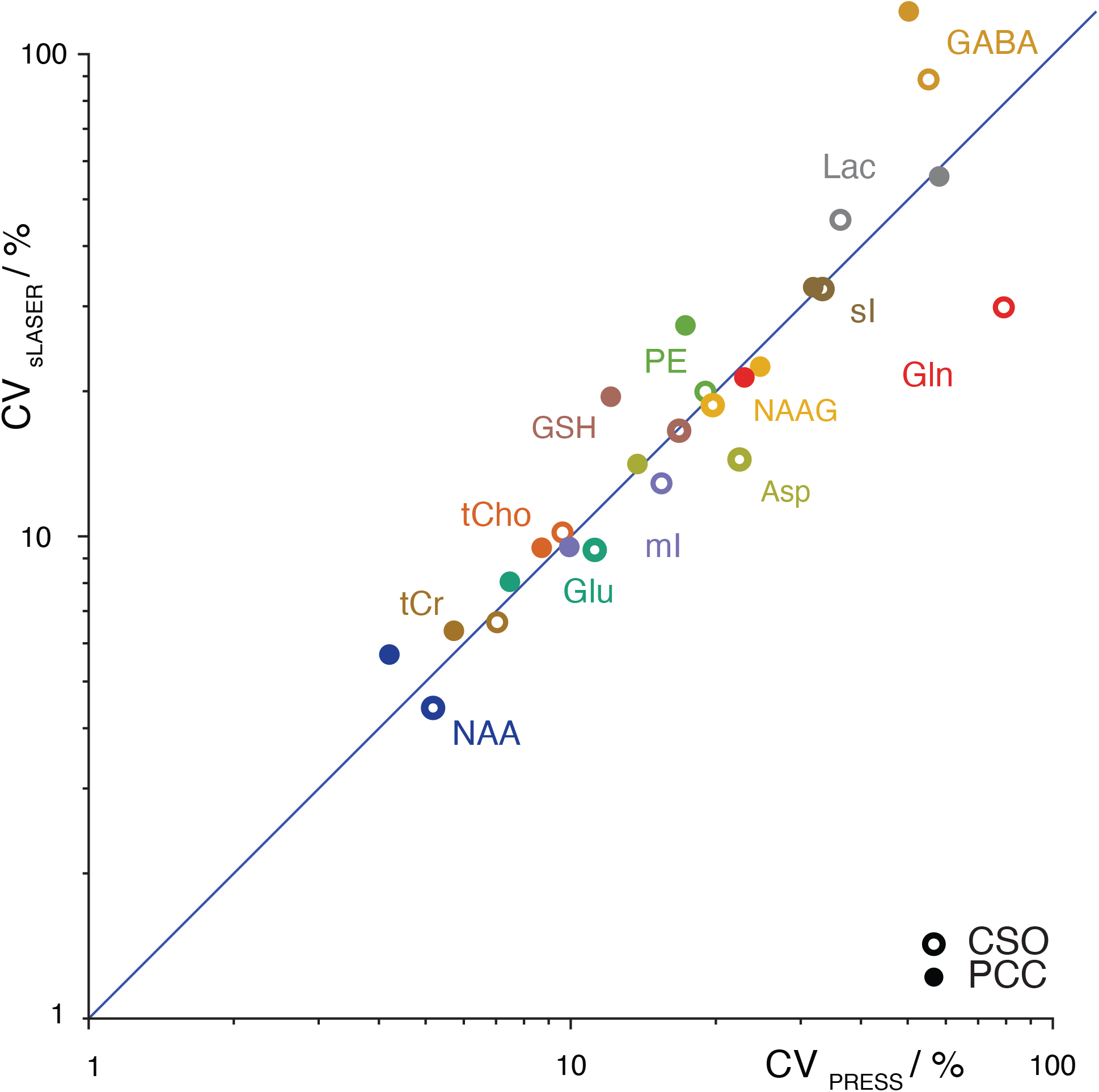
Coefficients of variation for PRESS and sLASER measurements plotted against each other on a logarithmic scale. CSO data are represented by open circles, and PCC by filled circles; colors indicate each metabolite.

## 4 DISCUSSION

Current MRS community consensus prefers sLASER over PRESS. This is mainly motivated by the reduced CSDA to minimize the location discrepancy between metabolites with different chemical shifts, which deteriorates as field strength increases. In this study, MRS data were acquired in a large healthy adult cohort using PRESS and sLASER. Our results suggested most metabolite measures are significantly correlated between PRESS and sLASER. Metabolite-age relationships were consistent between methods in CSO but not for 2 metabolites of interest (GSH and Glu) in PCC, in that *either* PRESS *or* sLASER (and *not* the other) suggested a significant age relationship. Differences in CVs between the methods were small in most metabolites. Overall, there is no strong evidence to support that either PRESS or sLASER outperforms the other for the analysis of brain metabolites using in vivo data acquired at 3T.

Our results indicate that spectral quality, including modeling residuals, is very similar for both methods. sLASER showed slightly better SNR and poorer linewidth, consistent with the increased sLASER signal mostly being located within the transition edges of the PRESS-localized voxel. The reduced CSDA of sLASER did not yield improved linewidth, presumably because the increased localization bandwidth does not give substantially better shim in the tCr-excited region.

PRESS and sLASER measurements were strongly correlated for tNAA, tCho, tCr, mI, and sI, and moderately correlated for most other metabolites. Even metabolites for which simple linear combination modeling of unedited spectra is questionable showed significant between-method correlations (e.g. Asp and PE in both regions and GSH in CSO). Where agreement between methods was surprisingly poor (e.g. Lac, Gln and GABA in CSO, GSH and NAAG in PCC), this could not be explained by differences in method performance, since the within-method variance in metabolite measures was similar.

Most of the metabolite concentration measurements were significantly different between PRESS and sLASER except tNAA and sI in PCC. Glu, Gln, tCr, mI, and Asp measurements were higher in PCC than in CSO for both PRESS and sLASER. NAAG, tCho and PE were higher in CSO. These results agree with previous literature examining gray versus white matter differences, except in the case of mI and PE (41). Concentration measurements of tCho, PE, GSH and GABA were higher in both CSO and PCC when using PRESS. Relaxation correction may contribute to these regional differences. Adiabatic pulses induce spin-locking in affected systems, extending the effective T_2_ relative to non-adiabatic sequences. Additionally, the three pairs of inversion pulses suppress diffusion-related coherence loss, as with a CPMG pulse train (42), lengthening apparent T_2_. While spectral differences resulting from scalar evolution are captured by simulation, metabolite relaxation corrections are often based on more-readily available non-adiabatic acquisitions, for example, using J-PRESS (43). Tissue-specific relaxation correction using a single set of relaxation times—as performed in Osprey—introduces metabolite- and region-specific offsets between PRESS and sLASER, which may account for some of the observed regional differences. If sLASER is to be fully adopted by the community a new set of sLASER-measured T_2_ reference values are probably required.

Relative to age, which is treated in this study as an external validation variable, there is no strong evidence to support sLASER outperforming PRESS at revealing biological metabolite relationships. Both PRESS and sLASER suggested significant age relationships in the same direction for tCho, tCr, mI, and GSH in CSO and for tCho, tCr, mI, and Asp in PCC. Interestingly, most age-metabolite correlation coefficients were slightly stronger for PRESS than sLASER. In PCC, there was a disagreement for GSH and Glu. PRESS data suggested a significant correlation for GSH which sLASER did not, and vice versa for Glu. However, overall, PRESS and sLASER produced very similar results in metabolite-age concentration relationships – for a majority of metabolites, they agreed either in observing a metabolite-age correlation or not observing any.

CVs for metabolite concentration measurements were similar for PRESS and sLASER. This is strong evidence for the consistency between methods in terms of their ability to acquire a similar dataset and quantify metabolites to a similar degree. The CVs of GABA and Gln were large (>50%) and PRESS and sLASER performed most differently for these metabolites (>30% difference). Quantification of these metabolites at 3T without editing is extremely challenging. GABA is often excluded from such analyses; these data suggest equal levels of caution are appropriate for quantification of Gln and Lac at 3T. Modeling of these lower-concentration metabolites is challenging, to the extent that the model returned zero values in multiple subjects. Lac can be measured with edited methods (as with GABA); Gln quantification is made challenging by overlapped Glu signals (they are often quantified in combination as Glx), although improved separation can be seen in J-resolved (44), TE-averaged (45) or edited sum spectra (46). High-concentration metabolites such as NAA, tCho, and tCr had low CVs and their differences between methods were also low (within 2% for PRESS and sLASER), indicating very consistent quantification between the two localization methods.

Minimizing CSDA at higher field strengths becomes crucial as CSDA increases with increasing field strength (47). Furthermore, at higher field strengths, the adiabatic pulses of the sLASER sequence prove increasingly beneficial as they are less impacted by inhomogeneous fields (12). It is important to emphasize that both brain regions studied here are ‘easier’ regions with average linewidths of 6Hz, indicating excellent shimming. It is possible that the benefits of improved sLASER localization would be more evident in more challenging brain regions with poorer shim (and particularly voxels adjacent to areas of bad shim), even at lower field strengths. However, there are also potential downsides to sLASER, including increased SAR from adiabatic (and more) pulses, increased minimum TE, and the larger number of potential coherence transfer pathways due to the larger number of pulses. Most of these must be suppressed by crusher gradients, to avoid an increased likelihood of exciting out-of-voxel artifacts in more challenging brain regions. In vivo evidence is required to establish whether sLASER is better than PRESS in such areas.

Procedures for data analysis in this study follow the community consensus by including an experimentally-derived MM basis function rather than using simulated parameterized Gaussian basis functions. A previous study investigated how MM modeling strategy impacted mean metabolite levels and found that incorporating a cohort-mean MM basis function in short-TE linear-combination modeling led to smaller model residuals and lower (i.e. better) Akaike information criterion values (33). Thus, given this evidence that an experimentally-derived MM basis function improves the overall estimates of metabolite measurements, we similarly implemented this modeling strategy in the present work.

## 5 CONCLUSION

Our results suggest that PRESS and sLASER perform similarly on brain metabolite measurements at 3T. High correlations and consistent measurements were observed in between-method and metabolite-age comparisons. CVs for metabolite concentration measurements were comparable between methods. These are important results as the current community consensus argues strongly for the use of sLASER, which is superior from the technical perspective, but lacks large-cohort in vivo evidence to support that it outperforms PRESS. Further experimental evidence is required to demonstrate that sLASER improves our ability to uncover new information about brain biochemistry; this study, the largest of its kind to-date, suggests that continued use of PRESS is entirely reasonable.

## Acknowledgement

This work was supported by NIH grants R01 EB016089, R01 EB023963, R21 AG060245, R00 AG062230, K99 DA051315 and P41 EB031771.

